# Exposure to nitric oxide drives transition to differential culturability in *Mycobacterium tuberculosis*

**DOI:** 10.1101/2021.09.28.462152

**Authors:** Sarah M. Glenn, Brindha Gap-Gaupool, Emily Milburn, Obolbek Turapov, Marialuisa Crosatti, Jennifer Hincks, Bradley Stewart, Joanna Bacon, Sharon L. Kendall, Martin I. Voskuil, Olga Riabova, Natalia Monakhova, Jeffrey Green, Simon J. Waddell, Vadim A. Makarov, Galina V. Mukamolova

## Abstract

During infection *Mycobacterium tuberculosis* (Mtb) forms differentially culturable (DC) subpopulations that are recalcitrant to treatment and undetectable using standard diagnostic tools. DC Mtb are revealed in liquid media, their revival is often stimulated by resuscitation-promoting factors (Rpfs), secreted peptidoglycan-remodelling enzymes, and prevented by Rpf inhibitors. Here we investigated the role of nitric oxide (NO) in generation of Rpf- dependent DC Mtb, using murine macrophage infection models and treatment with a synthetic NO donor (NOD). Mtb subpopulations were assessed by colony-forming unit counting on agar or by limiting dilution Most Probable Number assays in liquid media with or without Rpf inhibitor. Rpf-dependent DC Mtb were detected following infection of interferon-γ induced macrophages capable of producing NO, but not when iNOS was inactivated. NOD treatment also induced transition to the Rpf-dependent DC phenotype which was accompanied by global transcriptomic changes resulting in the dramatic down-regulation of *rpfA-E* gene expression. Furthermore, the DC phenotype was partially reverted by artificial over-expression of Rpfs. This study elucidates molecular mechanisms underlying the generation of DC Mtb, which are the dominant population recovered from clinical tuberculosis samples, with implications for improving both tuberculosis diagnostics and treatments.

## INTRODUCTION

*Mycobacterium tuberculosis* (Mtb) is the causative agent of tuberculosis (TB), which resulted in the death of 1.6 million people in 2021.^1^ Standard TB treatment lasts at least 6 months and significant resources are directed towards developing shorter and more effective treatments.^2^ One barrier to achieving this goal is the presence of Mtb populations that are often missed by standard diagnostic tests. These bacilli do not produce colonies on agar and only grow in liquid media, often requiring supplementation with culture supernatant (CSN)^3^ and are therefore known as differentially culturable (DC) Mtb.^4^ DC Mtb have also been referred to as non-culturable bacteria^5^ and differentially detectable bacteria.^6^ DC Mtb have higher tolerance to some anti-TB drugs^3,7^ and drug treatment increases the proportion of DC Mtb in patients.^3, 6, 8, 9, 10^ DC Mtb are also detectable in infected murine lungs and spleens^11, 12^ and show higher resistance to rifampicin.^11^ Thus, DC Mtb are believed to exist in a poorly characterized, non-replicating state, emergence from which requires liquid media or CSN for resuscitation and restoration of the ability to form colonies on solid media.

CSN contains resuscitation-promoting factor (Rpf) proteins, which are peptidoglycan-remodeling enzymes^13^ that can revive DC Mtb from the sputa of TB patients^3, 14^ and from animal lungs.^15^ Rpf inhibitors^16^ impair regrowth of DC mycobacteria from sputum^17^ and animal tissue.^15^ Furthermore, *ΔrpfB* and *ΔrpfAB* Mtb mutants exhibited reactivation defects in C57BL/6 mice treated with the iNOS inhibitor, aminoguanidine^18, 19^ or in iNOS-/- mutant mice^19^ suggesting an interplay between iNOS and Rpfs in controlling Mtb persistence *in vivo*. Whilst resuscitation of these Rpf-dependent DC Mtb has been linked to TB relapse in mice,^11^ the importance of this process in human TB remains to be established.

The molecular mechanism(s) underpinning the generation of DC Mtb is (are) currently unknown, although the *in vivo* environment has been proposed as a key factor.^15^ Nitric oxide (NO) is a well-characterized component of the innate immune system that is deployed to control intracellular pathogens, like Mtb.^20^ The inducible NO synthase (iNOS) of macrophages is critical for controlling Mtb in mice^21^ and humans.^22^ NO limits Mtb intracellular growth^23^ due to its direct antibacterial effects or by influencing the host cell mediated inflammatory response.^22^ Compounds that release NO (NO donors) produce a range of outcomes from bacteriostatic to bactericidal when added to Mtb cultures, dependent in part on the kinetics of NO liberation from the donor.^22^ Overall, results of NO exposure *in vitro* revealed relatively minor impacts on Mtb viability and cellular metabolism^22^ and have thus far failed to explain the significant effects of NO observed *in vivo*.^21, 23^

Here, it is shown that DC Mtb were formed during infection of murine macrophages, J774A.1 and CL57BL, stimulated with interferon gamma (IFN-γ), a known inducer of iNOS activity and NO production.^20, 23^ Resuscitation of these DC Mtb in liquid media supplemented with Rpf-containing culture supernatant (CSN), measured by limiting dilution Most Probable Number (MPN) assays, was abolished by an Rpf inhibitor, (3-nitro-4-thiocyanato-phenyl)- phenyl-methanone.^15, 16, 17^ These findings suggested that DC Mtb recovered from macrophages developed Rpf dependency. Rpf-dependent DC Mtb were not observed after infection of iNOS knockout murine macrophages or in cell lines treated with the iNOS inhibitor, aminoguanidine. To determine whether NO plays a central role in the generation of Rpf-dependent DC mycobacteria, cultures of Mtb and *Mycobacterium bovis* BCG (BCG) were treated with a new NO donor (NOD), which resulted in rapid generation of DC populations, resuscitation of which was abolished by the Rpf inhibitor. Transcriptomic analysis of NOD-treated cultures revealed widespread changes in gene expression, most importantly the down-regulation of *rpf* genes and identified possible regulatory networks underpinning this response. Furthermore, overexpression of *rpfD* or *rpfE*, using a tetracycline inducible system,^24^ partially reverted the Rpf-dependency phenotype and increased proportions of colony-forming bacteria on solid media after exposure to NOD. Thus, it was concluded that NO induces the Rpf-dependent DC phenotype of mycobacteria in both infected macrophages and in laboratory cultures.

## RESULTS

### Rpf-dependent DC Mtb are produced in IFN-γ activated murine macrophages

Mtb can survive and replicate in macrophages where they may be exposed to NO.^25^ Murine macrophage cell lines produce NO, especially after stimulation with IFN-γ.^23^ Two different murine macrophage cell lines (C57BL/6 and J774A.1) were used to investigate whether DC Mtb were formed intracellularly in response to NO exposure (Figure 1 and Figure S1). Resuscitation Indices (RIs) reflecting the difference between Mtb regrown in liquid media (MPN measurements) and Mtb producing colonies on solid agar plates (CFU measurements) were calculated using the following formula RI= Log_10_(MPN/ml) – Log_10_(CFU/ml).

**Figure 1.**
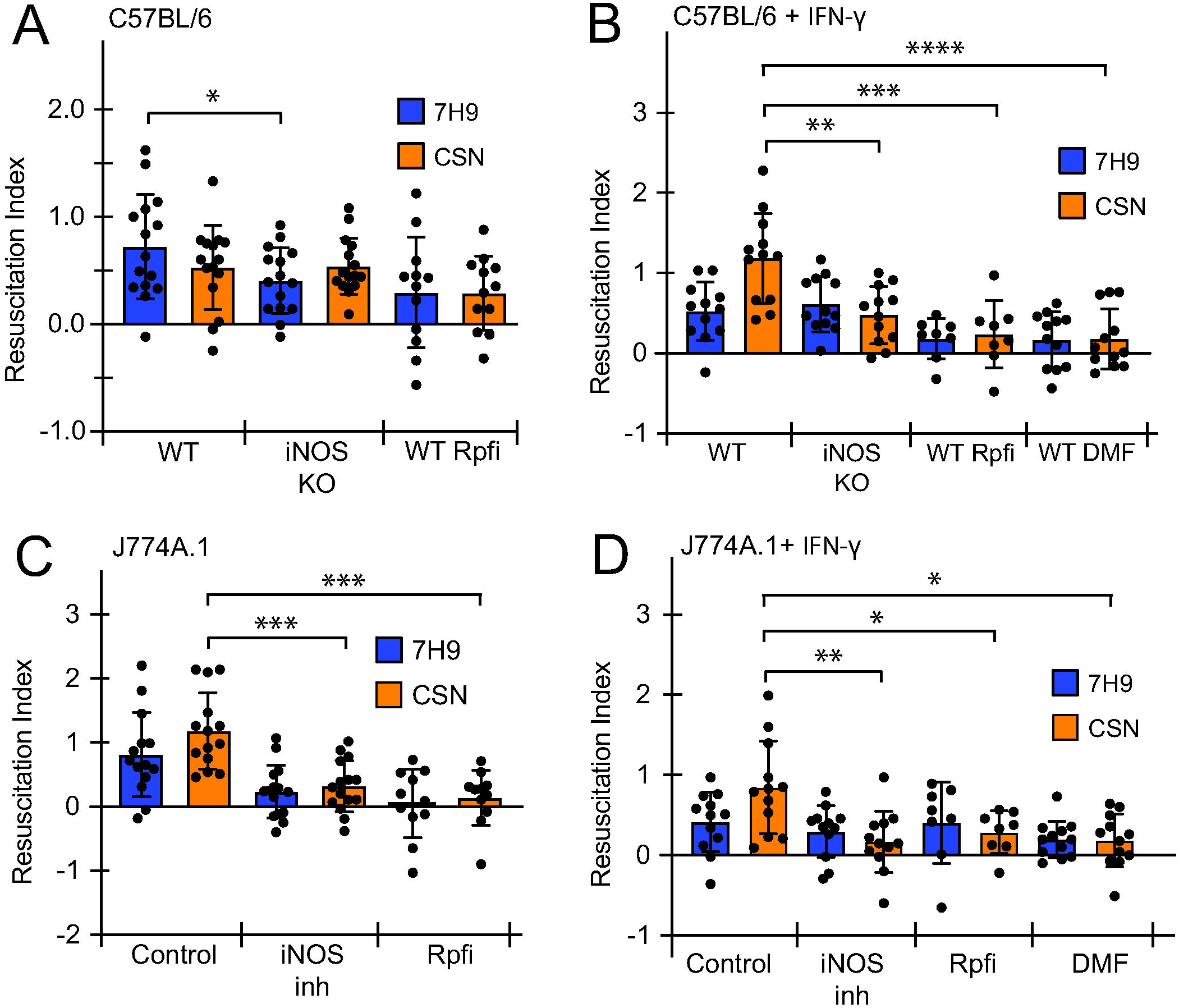
Rpf-dependent DC Mtb were formed in IFN-γ-stimulated murine macrophages with functional iNOS activity. Macrophages were infected with Mtb at MOI 1 for 24 hours prior CFU and MPN counting in 7H9 or CSN. The Rpf inhibitor (Rpfi) was added to resuscitation media as indicated. Data presented as RI values, RI = Log_10_MPN/ml - Log_10_CFU/ml. See Figure S1 for Mtb cell counts. (A, B) C57BL/6 wild type (WT) and iNOS knockout (iNOS KO); (A) untreated C57BL/6 or (B) IFN-γ treated C57BL/6. (C, D) J774A.1 cells; (C), untreated control or (D) IFN-γ treated cells. iNOS was inhibited by addition of aminoguanidine (iNOS inh). Data are means ± SEM from at least 12 biological replicates from 4 experiments. *p<0.05, **p=<0.01, *** p<0.001, ****p<0.0001 (unpaired t-test). Chemical concentrations used: IFN-γ, 1 ng/ml; aminoguanidine; 500 μM; DMF, 25 μM; Rpfi, 35 μM, (the concentrations of aminoguanidine and DMF were nontoxic for macrophages in accordance with previously published studies).^23, 61^

Figure 1A shows that C57BL/6 macrophages infected with Mtb for 24 hours had a DC Mtb population recoverable in 7H9 medium alone, as indicated by a resuscitation index (RI) of 0.7. This population was significantly lower in the infected C57BL/6 iNOS knockout macrophages (p<0.05). However, for both WT C57BL/6 macrophages and iNOS knockout C57BL/6 macrophages the RI values were not significantly greater when medium supplemented with CSN was used compared to 7H9 medium alone (Figure 1A). Moreover, application of an Rpf inhibitor (3-nitro-4-thiocyanato-phenyl)-phenyl-methanone),^16^ Rpfi, did not result in a statistically significant (p>0.05) decrease in RI values. These data suggested that the DC Mtb generated under these conditions were not dependent on exogenous Rpf proteins.

In the experiments described above, nitrite (a stable product of NO oxidation) was undetectable in the culture medium, indicating that NO production by C57BL/6 cells under these conditions was, at best, low. Therefore, the C57BL/6 macrophages were stimulated with IFN-γ resulting in detectable nitrite in the culture medium (2.02±0.18 µM, n=6 from two experiments). Now the DC Mtb population recovered in CSN (in the presence of exogenous Rpf) increased (RI=1.2) compared to both recovery in 7H9 medium (RI=0.52), Figure 1B, and to Mtb recovered from unstimulated C57BL/6 macrophages (Figure 1A). Addition of Rpfi dramatically decreased recovery of this population (RI=0.24). Furthermore, the DC Mtb population recovered in the presence of CSN from the IFN-γ stimulated iNOS knockout macrophages was significantly smaller in (p<0.01) than that from stimulated WT macrophages (Figure 1B) and resuscitation in both 7H9 and CSN supplemented media was prevented by addition of an Rpfi (Figure 1B). These data show that Rpf-dependent DC Mtb were formed in IFNγ-stimulated, NO-producing WT C57BL/6 macrophages.

Previous studies established that DC Mtb were eliminated from infected murine organs by treatment with the anti-inflammatory compound dimethyl fumarate (DMF).^12^ This was also the case for infected IFNγ-stimulated WT C57BL/6 macrophages (p<0.0001), where the Mtb RI was reduced to 0.2 (Figure 1B).

The NO-dependent generation of DC Mtb in macrophages was next investigated using a second cell line, J774A.1. After 24 hours of infection, DC Mtb were recovered from unstimulated macrophages, but not from J774A.1 cells with chemically inactivated iNOS, using the murine iNOS inhibitor aminoguanidine (Figure 1C). DC Mtb could be recovered in 7H9 medium (RI=0.8), and addition of CSN improved resuscitation (RI=1.2), which was abolished by the addition of the Rpf inhibitor (RI<0.2). In IFNγ-stimulated macrophages the DC Mtb population was only recoverable in CSN-containing medium (RI=0.85), and significant DC Mtb populations were not produced after treatment with aminoguanidine (RI=0.17), or the Rpf inhibitor (RI=0.29) or DMF (RI=0.27) (Figure 1D). Collectively, our data suggest that generation of Rpf-dependent DC Mtb in macrophages can be triggered by NO and is prevented by iNOS inhibitors or the anti-inflammatory drug, DMF.

### Treatment with a novel nitric oxide donor induces Rpf dependency in Mtb and BCG

We next investigated whether DC mycobacteria could be induced by exposure to NO in culture. A new NO donor (NOD), 3-cyano-5-nitropyridin-2-yl diethyldithiocarbamate (Figure 2A) for sustained intracellular NO exposure and a structurally related control compound (CC), 3-cyano-4,6-dimethyl-5-nitropyridin-2-yl piperidine-1-carbodithioate (Figure 2B) were designed and synthesized. Accumulation of NO in culture medium was assessed by measurement of nitrite, a stable oxidation product of NO;^26^ intracellular NO was detected by staining with DAF-FM diacetate.^27^ NOD did not release NO spontaneously in 7H9 medium, where pH was 6.8 (Figure S2A); however, a rapid increase of fluorescence was observed (within 1 hour) in NOD-treated BCG, but not in CC-treated mycobacteria (Figure 3A, S2B). Moreover, a significant DAF-FM diacetate positive BCG population could be detected after 24 hours of exposure to NOD; this was not the case for CC-treated, untreated, or heat-killed BCG treated with NOD (Figure 3B). To confirm our findings, we measured nitrite concentrations in bacterial lysates obtained from treated and untreated BCG cultures (Figure 3C). NOD treatment resulted in significant increase of nitrite within 1 hour as compared with CC-treated and untreated BCG cultures (p<0.001, one way ANOVA). Furthermore, more than 50% of cells in NOD- and CC-treated samples were propidium iodide negative and SYTO 9 positive, and therefore likely remained viable (Figure S2C).

**Figure 2.**
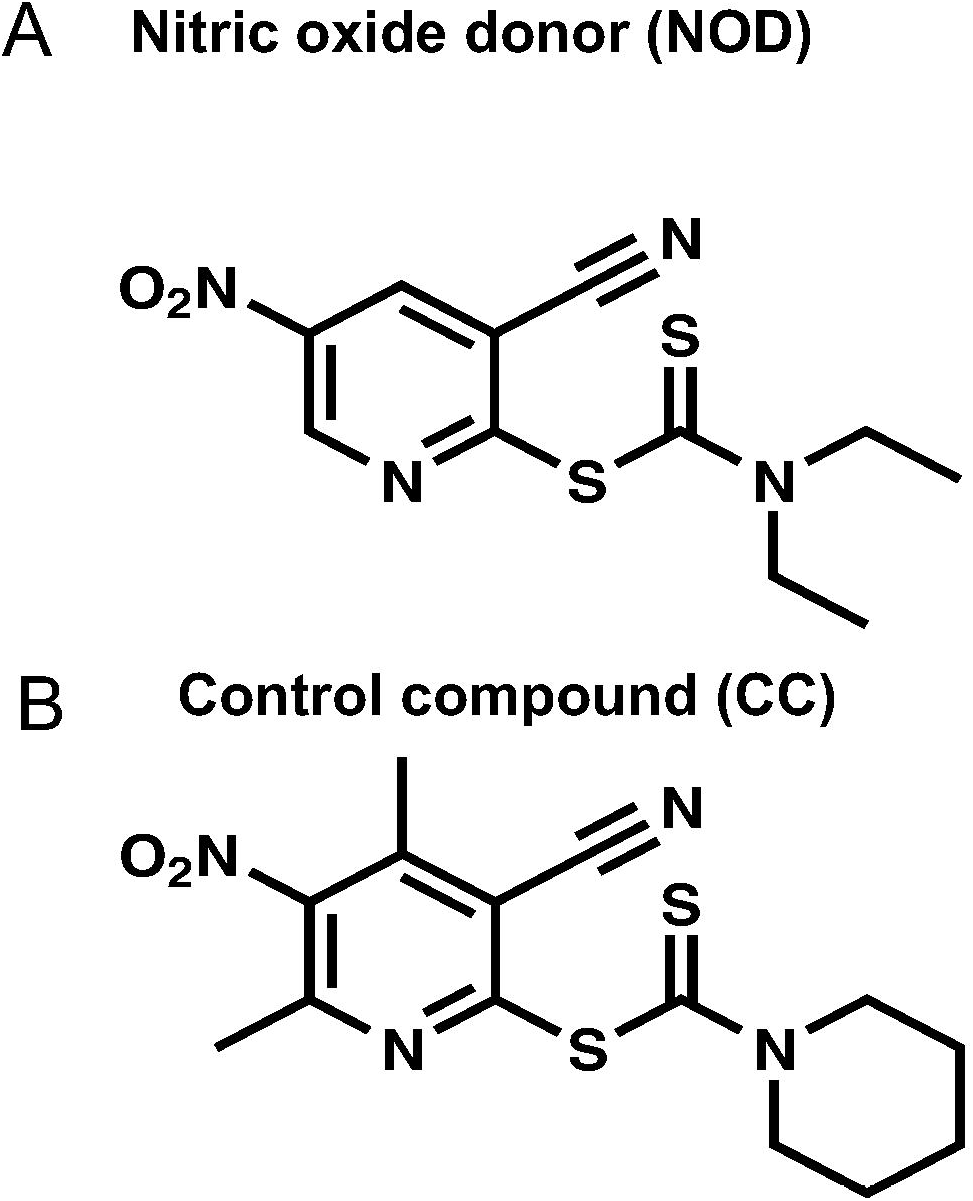
Chemical structures of (A) the NO donor, 3-cyano-5-nitropyridin-2-yl diethyldithiocarbamate, NOD and (B) the control compound, 6-dimethyl-5-nitropyridin-2-yl piperidine-1-carbodithioate, CC.

**Figure 3.**
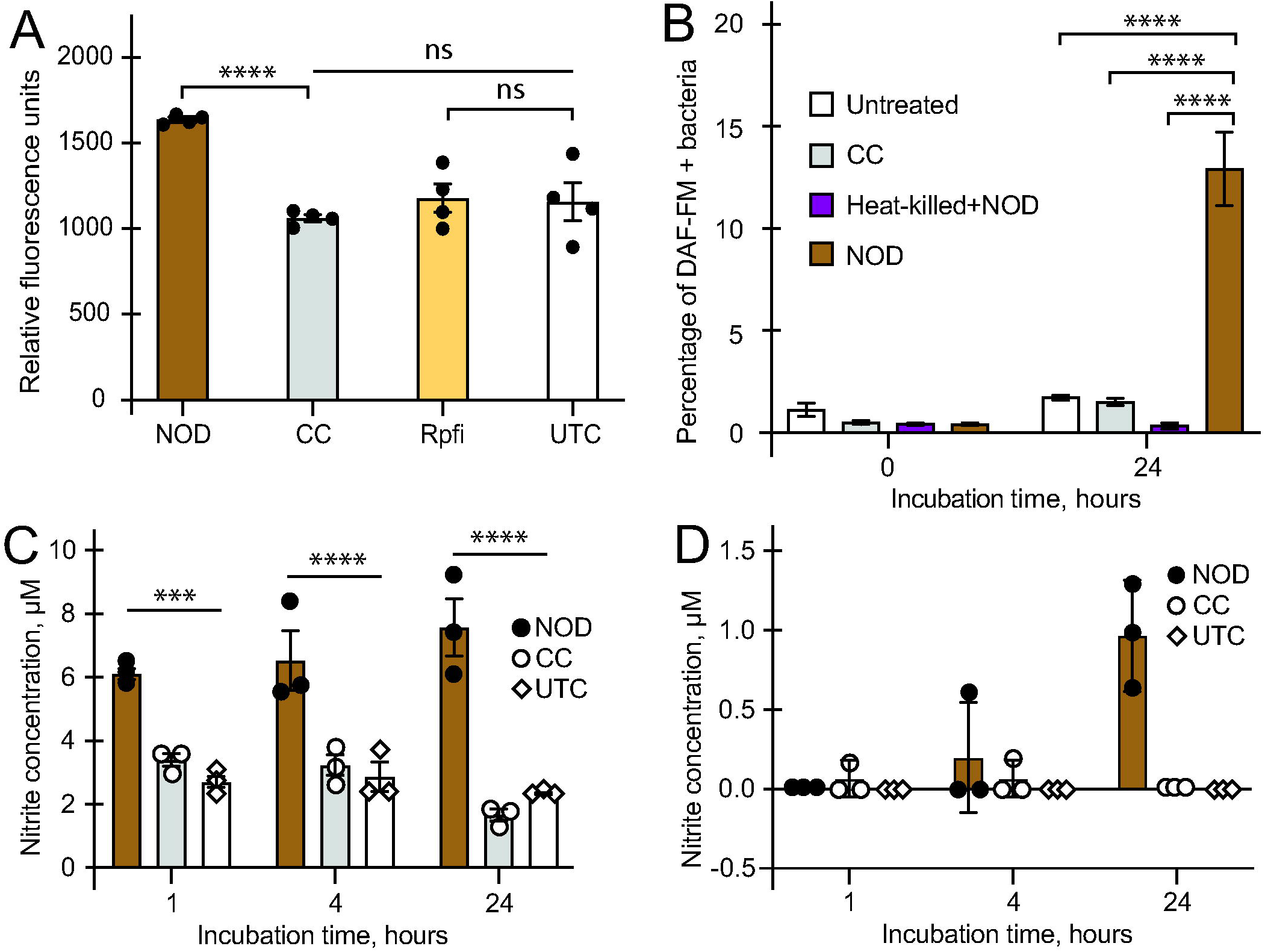
NOD enabled intracellular NO release. (A) Detection of NO in BCG using DAF-FM diacetate and fluorescence measurement. Bacteria were either untreated (UTC) or incubated with NOD, CC, or Rpf inhibitor (Rpfi) for 1 hour. Data are means ± SEM for 4 biological replicates from one representative experiment ****p<0.0001, unpaired t-test. (B) Detection of DAF-FM diacetate positive cells in BCG cultures treated with either 100 µM NOD or CC for 24 hours using flow cytometry. As well as untreated cultures (Untreated) the control experiments included heat-treated BCG mixed with 100 µM NOD (Heat-killed + NOD). ****p<0.0001 (unpaired t-test). (C, D) Assessment of nitrite concentration in lysates (C) or spent media (D) prepared from BCG treated with either NOD or CC for 24 hours. ****p<0.0001, one way ANOVA. Chemical concentrations used: NOD or CC, 100 µM (A, B) and 200 µM (C, D); Rpfi, 35 µM; DAF-FM diacetate, 10 µM. (B-D) Data are means ± SEM for three biological replicates from one experiment.

These observations suggested that NOD entered live BCG, releasing measurable levels of NO without causing bacterial cell lysis. Therefore, NOD was deemed suitable for testing the effects of NO exposure on growth phenotypes using CC as a control.

While incubation of Mtb with 100 µM CC for 24 hours had no effect on cell counts in liquid or on solid media (Figure S3A), exposure to NOD resulted in a 1000-fold reduction in CFU counts (Figure 4A). In liquid 7H9 medium the recovered cell counts increased ∼100-fold and supplementation with CSN further increased this number >10-fold (Figure 4A). More detailed analysis revealed a gradual decline of CFU counts in cultures of NOD-treated Mtb over 48 hours (Figure S3B). This suggested that, like Mtb from clinical TB samples, most of the NOD-treated cells were DC Mtb.^3,4^ Similar results were obtained with BCG (Figure 4B). BCG cultures had lower CFU counts after 24 hours compared to Mtb (Figure 4B) and resuscitation indices (difference between CFU and MPN counts) were slightly higher in BCG compared with Mtb cultures (Figure 4C). As might be expected, differences were observed between experiments carried out at different times with independent cultures; however, the RI values for NOD-treated mycobacteria measured in the presence of CSN were always greater than those obtained with unsupplemented 7H9 medium (Figure 4C). Thus, these results revealed that NOD treatment of both Mtb and BCG led to formation of DC bacteria.

**Figure 4.**
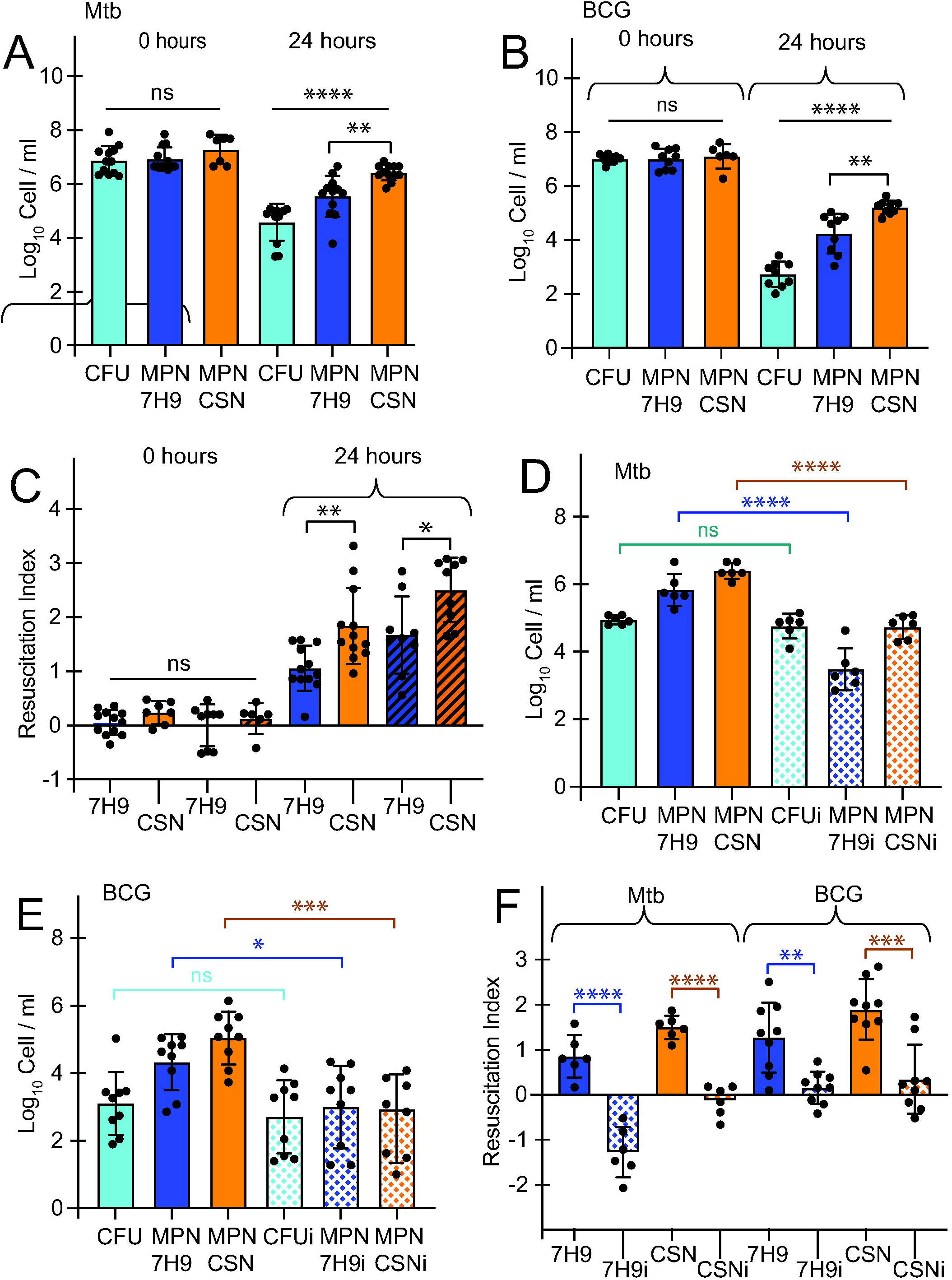
NOD treatment induced Rpf-dependent mycobacteria *in vitro*. (A-E) Mtb or BCG were treated with 100 µM NO donor (ND) for 24 hours at 37°C without shaking. CFU, MPN 7H9 (MPN in 7H9 liquid medium) and MPN_CSN (MPN in CSN) counts for Mtb (A, E) and BCG (B, D) were determined. (C) Resuscitation indices (RI) were calculated for Mtb (filled bars) and BCG (hatched bars). (F) RI were calculated for Mtb and BCG samples resuscitated with (dotted bars) or without Rpfi. (D-F) Rpfi (35 µM) was added to resuscitation media, 7H9 or CSN (“i” - resuscitation in the presence of Rpfi) and cell counts determined for Mtb (D) or BCG (E). (A, B, D, E) ****p<0.0001, one-way ANOVA; (C, F) *p<0.05, **p<0.01, unpaired t-test. Data are means ± SEM for at least six replicates from at least two experiments. RI = Log_10_MPN/ml - Log_10_CFU/ml.

The addition of the Rpfi to resuscitation media (7H9 or CSN), abolished the resuscitation of NOD-treated Mtb (Figure 4D) or BCG (Figure 4E). Figure 4F shows RI values calculated for NOD-treated Mtb and BCG resuscitated in media with or without Rpfi and confirms that Rpfi had a similar effect on resuscitation of both mycobacteria. In accordance with previous findings,^16^ the Rpfi had no effect on actively growing Mtb (Figure S4). Rpfi also did not increase DAF-FM diacetate fluorescence (Figure 3A).

### NOD treatment produced a broad transcriptional response and down-regulated expression of *rpf* genes

To investigate molecular mechanisms controlling formation of DC Mtb in response to treatment with NOD, transcript-profiling experiments were conducted. Cultures were sampled 4 hours after exposure to either NOD or CC. At this time, for NOD-treated cultures, the CFU count was decreasing but the MPN_CSN value was not, suggesting that the transition to the DC state had begun (Figure S3B). Compared to treatment with CC, 640 (407 induced, 233 repressed) genes were differentially expressed, fold change >2, p<5×10^-5^ (Figure 5A, Dataset 1). The changes in gene expression spanned a wide range of functional categories (Figure 5B). The DosR regulon that has been previously shown to react to hypoxia and NO; signals associated with non-replicating states of Mtb^28, 29, 30, 31^ was not induced after NOD treatment (Dataset 1). Moreover, the Mtb *dosR* deletion mutant, a complemented strain, and the parent Mtb with NOD for 24 hours resulted in similar numbers of DC Mtb; all strains were equally resuscitated in CSN-supplemented medium (Figure S5A).

**Figure 5.**
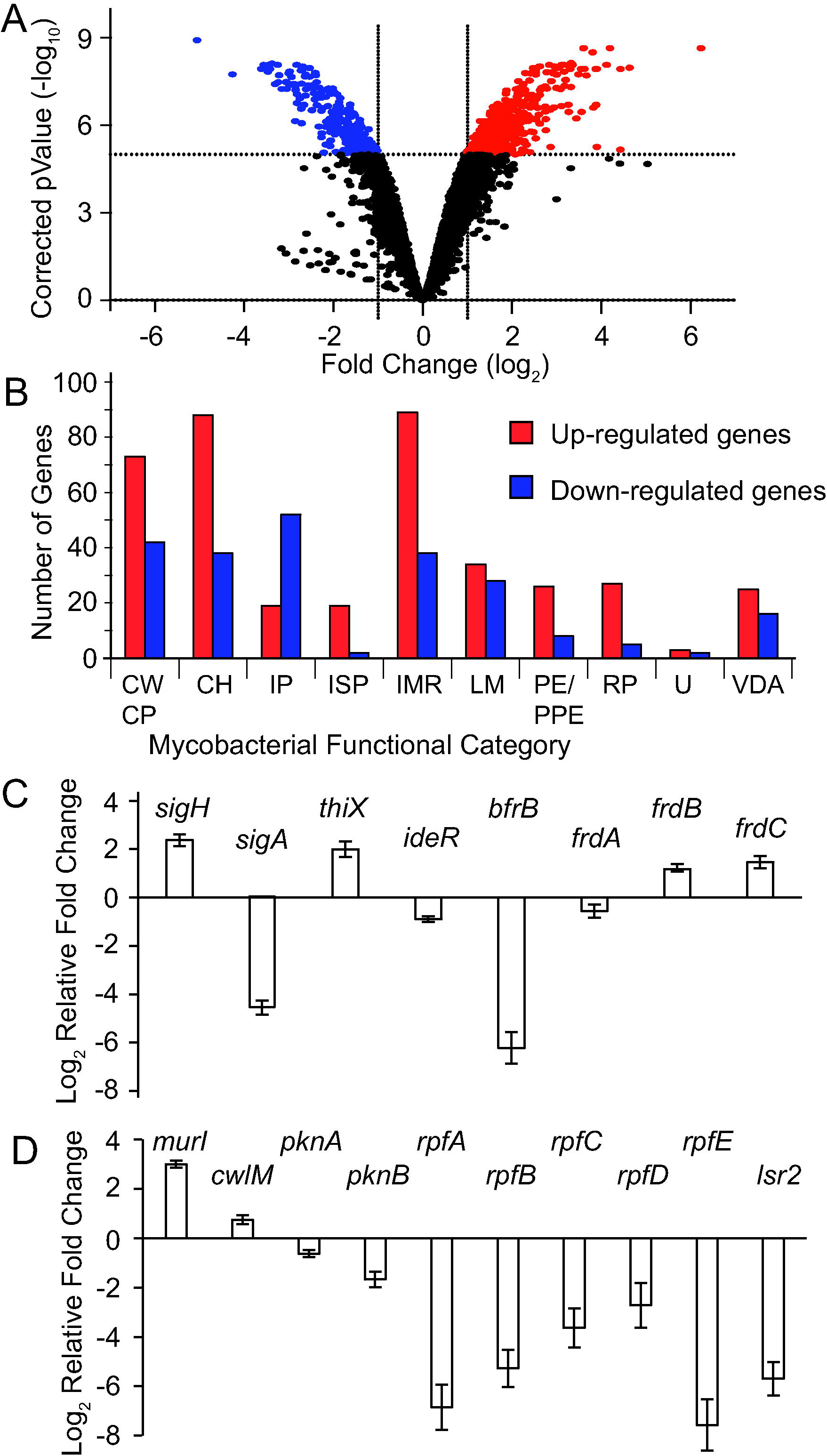
Mtb exposure to NOD for 4 hours led to activation of several regulatory pathways. (A) Volcano plot showing 407 genes significantly induced (red) and 233 genes repressed (blue) by NOD compared to CC at 4 hours. Significantly differentially expressed genes (DEG) were identified using a moderated t-test (p<5×10^-5^ with Benjamini and Hochberg multiple testing correction) and fold change >2.0. (B) Distribution of DEG from functional categories.^62^ CWCP – cell wall and cell processes; CH – conserved hypothetical; IP – information pathways; ISP – insertion sequences and phages; IMR – intermediary metabolism and respiration; LM – lipid metabolism; RP – regulatory proteins; U – unknown; VDA – virulence, detoxification, adaptation. (C, D) Expression of selected genes was confirmed by RT-qPCR, normalized to 16S rRNA transcript abundance. Log_2_ relative fold difference was calculated. Data are means ± SEM for three independent cultures done in technical duplicates (n=6).

Several transcriptional changes indicated adaptation to impaired aerobic respiration (Dataset1). These included up-regulation of fumarate reductase (*frd*) genes (Figure 5C), suggesting utilization of fumarate as a terminal electron acceptor to maintain redox balance and membrane potential,^32^ and the up-regulation of the *pta-ackA* operon, coding for phosphotransacetylase and acetate kinase, indicating increased reliance on substrate-level phosphorylation for ATP generation (Dataset 1). We note that expression of *icl1*, coding for isocitrate lyase, an enzyme of the glyoxylate cycle that is critical for survival under hypoxia,^33^ was down-regulated (6.15-fold) in response to NOD treatment (Dataset 1). Like the absence of a DosR regulon response, the down-regulation of *icl1* contrasts with several other studies of NO-treated Mtb in which *icl1* was up-regulated.^30, 31^ Taken together, these differences in gene expression suggest that NOD induces distinct physiological adaptations compared to other models. However, we note that previous studies showing strong induction of the DosR regulon by NO used well-aerated cultures, conditions that likely result in initial suppression of the DosR regulon. In contrast, here to better mimic the *in vivo* environment, we used static Mtb cultures in which the DosR regulon is likely to be already, at least partially, induced.^29^

The 407 up-regulated genes were analyzed using the Transcription Factor Over Expression (TFOE) tool^34^ to identify transcription regulators that mediate the response to NOD treatment. Seven transcription regulators passed the significance threshold (TFOE output: significance of enrichment of their regulons p<5×10^-5^), including three alternative sigma factors (SigH, SigF and SigK) and four transcription regulators (Rv0047, WhiB2, Rv2175 and FurA, Dataset 2). SigH and SigF were also prominent (significance p<0.05) in the ChIP-seq output (Dataset 2). SigH coordinates the response to oxidative, nitrosative and heat stresses in Mtb, and *sigH* expression is induced in macrophages (reviewed by Manganelli).^35^ Additionally, SigH is activated by oxidation of its anti-sigma factor RshA, which can be triggered by NO. SigF is associated with cell envelope integrity and is controlled by anti-sigma factor agonists, RsfA and RspB, which sequester the anti-sigma factor RsbW, and also by the anti-sigma factor Rv1364c. SigK is likely involved in maintaining the redox state of the mycobacterial periplasm.^40^

The involvement of SigH, SigF and SigK in the response to NOD suggests that Mtb was exposed to oxidative, nitrosative and cell wall stresses during NOD-induced transition to Rpf- dependency. Accordingly, expression of antioxidant genes, such as the SigH/AosR-activated non-canonical cysteine biosynthetic genes (*mec-cysO-cysM*), ergothioneine biosynthesis genes (*rv3700c* and *rv3704c*), sulfate-containing compound ABC transporter (*cysA1*), sulfate adenyltransferase (*cysDN*), methionine sulfoxide reductases (*msrA* and *msrB*), rhodanese domain protein (*rv1674c*), thioredoxin reductase and thioredoxin proteins (*trxB2-C*, *thiX*, *trxB1*) were up-regulated. These antioxidant responses are consistent with adaptation to manage NO-mediated oxidative damage.

As a radical, NO reacts with transition metals such as iron resulting in damage to iron-sulfur clusters and perturbation of redox balance.^36^ Disrupted iron homeostasis was suggested by the up-regulation of mycobactin biosynthetic genes (*mbtE*, *mbtG* and *mbtH*) and ESX-3 genes required for siderophore-mediated iron uptake (*rv0282-rv0292*), and down-regulation of *ideR* repressor and iron storage systems, *bfrB* (Dataset 1, Figure 5C).

The up-regulation of *murI,* coding for glutamate racemase, and *cwlM*, an essential regulator of peptidoglycan synthesis,^37^ indicated that remodeling of the cell wall was involved in the transition to Rpf-dependency (Dataset 1, Figure 5D). The up-regulation of *murI* is likely to be mediated by SigH (overexpression of *sigH* increased *murI* expression)^35^ and expression of *cwlM* was previously shown to be up-regulated by treatment with spermine NONOate.^38^

Consistent with the observation that NOD-treatment resulted in the appearance of Rpf- dependent mycobacteria, three *rpf* genes were down-regulated (*rpfA, rpfB* and *rpfE*; 9.5-, 3.9- and 33.2-fold, respectively) in the transcriptomic dataset (Dataset 1). This response was confirmed by RT-qPCR experiments, which showed that all five *rpf* genes were down-regulated in NOD-treated Mtb (Figure 5D). Expression of Rpf coding genes is tightly controlled by multiple regulators,^39^ including Lsr2^40^ which was also down-regulated in NOD-treated Mtb.

### Ectopic expression of *rpf* genes impairs NOD-induced generation of DC Mtb

Transcript profiling and RT-qPCR analyses suggested that exposure of Mtb to NOD resulted in lower expression of all five *rpf* genes. Our proteomics analysis showed that RpfA, RpfB and RpfD were present in the Mtb CSN used to reveal the presence of DC Mtb in macrophages and in cultures treated with NOD (Table 1). Furthermore, RpfA, RpfC and RpfE proteins have been previously detected in CSN.^41^ Thus, it was suggested that down-regulation of *rpf* genes in response to NO was important for the generation of Rpf-dependent DC mycobacteria. To test this hypothesis we constructed Mtb strains in which expression of either *rpfD* or *rpfE* was placed under the control of a tetracycline-regulated promoter, removing them from their native regulatory networks to permit sustained expression in the presence of NOD by addition of the inducer, anhydrotetracycline. The *rpfD* and *rpfE* genes were chosen for these experiments because reliable over-expression of both had been confirmed by qPCR and western blot experiments in previous studies.^24^ Furthermore, RpfD has been detected in CSN, and recombinant RpfE resuscitated DCB from sputum.^3, 14^ Ectopic expression of *rpfD* or *rpfE* resulted in an ∼100-fold increase in CFU counts compared with the empty vector control after NOD treatment (Figure S5B), demonstrating partial restoration of Mtb culturability. Together, these data support an important role for NO-mediated down-regulation of *rpf* expression for the formation of DC Mtb, consistent with their Rpf-dependency phenotype.

**Table 1.**
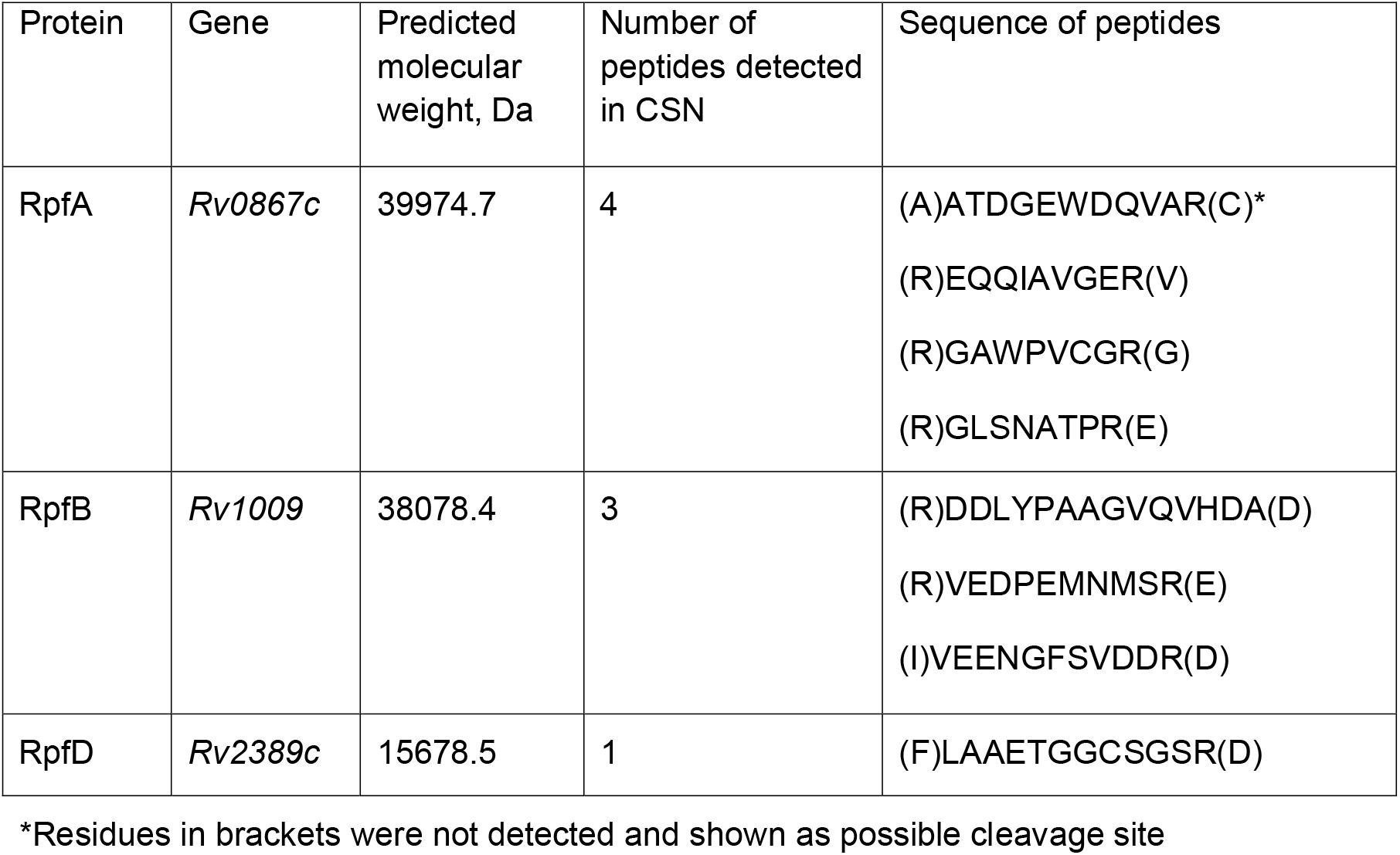
Rpf peptides detected in Mtb CSN.

## DISCUSSION

During infection pathogenic mycobacteria produce heterogeneous populations that differ in their metabolic activity, growth characteristics and tolerance to antimicrobials (reviewed by Chung *et al*.).^42^ Amongst these populations, differentially culturable (DC) mycobacteria are of particular interest, because they are difficult to detect and eradicate, and frequently represent the dominant Mtb population in clinical TB samples.^3, 4, 43, 44, 45^ By genetic or chemical manipulation of NO production by two macrophage cell lines, along with application of exogenous Rpf proteins (as CSN) and use of a specific Rpf inhibitor, we have shown that NO induces the generation of Rpf-dependent DC Mtb in macrophages. These observations suggest that host NO triggers this adaptive response that might be important for TB pathogenesis.

The observation that NO plays a major role in the formation of Rpf-dependent DC Mtb in macrophages prompted experiments to learn more about the mechanisms underpinning this response. In previous work, various strategies have been developed, including prolonged incubation in stationary phase,^5^ gradual acidification,^46^ alteration of sodium-potassium ratio,^47^ treatment with antimicrobials^11, 24, 48^ or inducers of oxidative stress,^49^ to study the formation of DC mycobacteria *in vitro*. Here we describe a model system for generating DC mycobacteria (both Mtb and BCG) based on a new NO donor (3-cyano-5-nitropyridin-2-yl diethyldithiocarbamate) that simulates the *in vivo* environment by generating heterogeneous mycobacterial populations, thereby permitting investigation of the physiologically distinct states produced during the pathogenesis of TB. Our data suggest that NOD treatment results in release of NO in living bacteria. Establishing the precise mechanism of NO release from NOD in mycobacterial cells, including any cellular factors involved, requires further investigation. Nevertheless, a low active concentration of NOD in comparison with other NO donors, and direct detection of NO in NOD-treated mycobacteria even after 24 hours of incubation, and the availability of a structurally similar control compound, makes NOD advantageous for investigation of bacterial physiology.

In our NOD-induced system, transition to the DC state was likely mediated by two alternative sigma factors, SigH and SigF and their regulons, previously implicated in adaptation to redox stress and respiration-inhibitory conditions.^35^ Overall, the transcriptomic signature of NOD-treated Mtb was associated with inhibition of cell division, repair of NO-induced damage, and metabolic reprogramming to restore redox balance and membrane potential. Whilst there was overlap between the transcriptional response to NOD treatment and those reported for other NO donors, there were several distinctive features associated with NOD-mediated formation of DCB, including the absence of a DosR response and the down-regulation of *icl1*. These changes were also observed in Mtb surviving prolonged multi-drug treatment in mice for 28 days.^50^ Remarkably, 4 out of 5 *rpf* genes (*rpfB-E*) were also downregulated in this model, while *murI* was up-regulated similarly to the NOD treated Mtb. We have previously shown that DC Mtb were the dominant population in mice treated with a combination of rifampicin, isoniazid and pyrazinamide for 28 days,^12^ therefore these transcriptomic adaptations might be attributed to DC Mtb. Interestingly, the transcriptional response to NOD treatment included genes from the recently identified signature of differentially detectable (DD) Mtb from sputum samples.^51^ In particular, 5 genes, *icl1*, *ppsA*, *hspX*, *rv1738* and *pks15*, were significantly down-regulated in sputum Mtb DD^51^ and NOD-treated Mtb (Table S2). Furthermore, *icl1* was down-regulated in another *in vitro* model for DD Mtb (the PBS-RIF model) and increased numbers of DD bacilli were formed by an *icl1* mutant.^49^ Together these observations suggest that down-regulation of *icl1* and the glyoxylate cycle might be an important regulatory and metabolic adaptation in response to multiple stimuli for the formation of DCB both *in vitro* and *in vivo*.

Although the deployment of stress response systems and changes in metabolic mode are likely to be important adaptations in the transition to the DC state, one of the most striking transcriptional changes observed was the down-regulation of all five *rpf* genes in Mtb cultures treated with NOD. Whilst there is an ongoing debate concerning the importance of NO in controlling Mtb infection in humans, even though NO is detected in TB patients,^52^ our data suggest that NO contributes to the generation of DC Mtb by down-regulation of *rpfA-E* expression. Rpfs are cell wall cleaving enzymes and their activities under conditions when the peptidoglycan producing machinery is likely to be damaged could be fatal for mycobacteria as a result of uncontrolled cell lysis and death. Accordingly, expression of Rpf coding genes is controlled by multiple regulators^39^ and involves post-translation regulations^53, 54, 55^ (Figure S6); however, at this time which of these contributes to down-regulation of these genes upon exposure to NO is unknown. Nevertheless, based on the evidence presented here, the simplest explanation for the observed Rpf-dependency of DC Mtb recovered from macrophages or after treatment with NOD is that exposure to NO down-regulates *rpf* gene expression such that mycobacteria fail to initiate Rpf production when grown in standard media. This means that emergence from the DC state requires exogenous supplementation with recombinant Rpf or Rpf-containing CSN to restart growth and peptidoglycan biosynthesis. This failure to initiate *rpf* expression could be caused directly by the action of NO on transcriptional regulatory networks or indirectly, via metabolic reprogramming in response to NO. In addition, NO also has the potential to directly damage Rpf proteins, which require a disulfide bond for correct folding and activity. The precise mechanism of Rpf-mediated resuscitation is currently unknown, and we cannot exclude that the Rpf effect is indirect and that resuscitation involves products of peptidoglycan degradation or other unknown factors. For example, in addition to Rpf proteins, various other molecules, including cAMP, muropeptides, phospholipids, and peptides derived from Rv1174c have been proposed to resuscitate DC mycobacteria (summarized by Dartois *et al*.).^56^ However, none of these additional factors have been systematically validated for resuscitation of DC mycobacteria from sputum in the way that Rpf proteins have. Indeed, CSN obtained from the quintuple *rpf* deletion mutant had variable resuscitation effects.^3,4^ Moreover, there is currently no published data on the composition of CSN from the quintuple *rpf* deletion mutant to suggest the identities of any alternative resuscitation factors. These observations serve to highlight the complexity of Mtb resuscitation and the need for further research to uncover the roles of any Rpf-independent resuscitation factors in that process. Rpf-dependent and Rpf-independent DC Mtb may represent different subpopulations and their formation might be induced by different factors.

Overall, our findings suggest that differential culturability is a survival mechanism that allows mycobacteria to cope with prolonged host-imposed stresses, and that this has implications for the development of better TB therapies and diagnostic tools. Exposure to various levels of NO drives Mtb population heterogeneity in the infected host (Figure 6). These heterogeneous populations have different growth requirements, respond differently to drugs and may induce differential immune responses. Here we show that NO down-regulates the expression of all *rpf* genes and generates Rpf-dependent DC Mtb intracellularly in macrophages and in a new laboratory model for studying the transition to differential culturability. Thus, our work provides a foundation for further investigations to understand the molecular mechanisms that underpin this fundamental adaptive process and elucidate the precise role of DC Mtb in TB pathogenesis and treatment outcomes.

**Figure 6.**
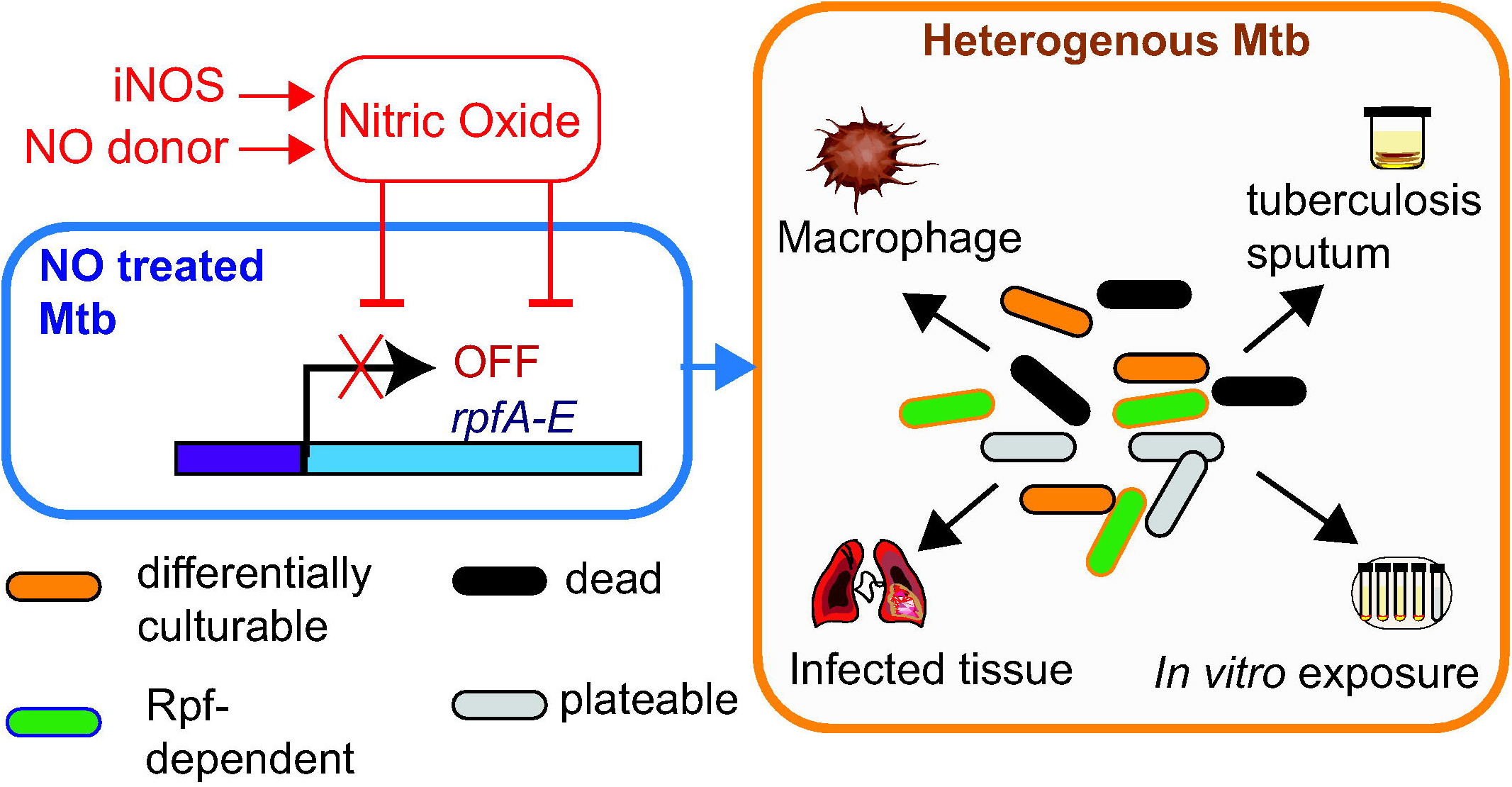
NO-mediated generation of Rpf-dependent DC Mtb. NO exposure *in vivo* and *in vitro* shuts down expression of *rpfA, rpfB, rpfC, rpfD* and *rpfE*. This leads to generation of heterogenous Mtb subpopulations including Rpf-dependent Mtb. This population heterogeneity mimics *in vivo* observations where NO is produced by the inducible NO synthase (iNOS) and drives formation of Rpf-dependent and differentially culturable Mtb that can be detected in infected tissue and sputum samples.

## MATERIALS AND METHODS

### Materials and Strains

Liquid cultures were grown at 37°C with shaking (100 rpm) for Mtb and static for *M. bovis* BCG Glaxo in Middlebrook 7H9 broth, supplemented with 0.2% (v/v) glycerol, 10% (v/v) albumin-dextrose-catalase (ADC), and 0.05% (w/v) Tween 80 (hereafter 7H9 medium). Culture supernatants (CSN) were prepared from exponentially growing cultures (OD_580nm_ of 0.6-0.8) and sterilized by double filtration (0.22 µm filters). For MPN assays, 7H9 medium was supplemented with 50% sterile CSN . The Rpf inhibitor (3-nitro-4-thiocyanato-phenyl)-phenyl-methanone^16^) was added to resuscitation media to a final concentration of 35 µM; a concentration based on previously published data.^15, 16, 17^ Mtb strains for overexpression *rpfD* and *rpfE* were obtained by electroporation of previously generated constructs;^24^ expression of *rpf* genes was induced by addition of anhydrotetracycline (20 ng/ml). The *dosR* deletion mutant and complement strain were previously described.^57^ Mtb stocks for macrophage infection were prepared from exponentially growing cultures (OD_580nm_ ∼0.4-0.5). J774A.1 (ATCC® TIB-67™), C57BL/6 wild type and iNOS knockout mutant (Kerafast) cell lines were grown at 37°C with 5% CO_2_ in Dulbecco’s Modified Eagle’s medium (DMEM) supplemented with 10% (v/v) heat inactivated fetal bovine serum. Interferon-γ (IFN-γ), dimethyl fumarate (DMF) and aminoguanidine and were added to final concentrations of 1 ng/ml, 25 µM and 500 µM respectively.

### Macrophage Infection

Murine macrophages (C57BL/6 or J774A.1) were seeded at 2×10^5^ cells/ml in 24 well plates (Greiner Bio-One). IFN-γ was added 24 hours prior to infection, aminoguanidine or DMF were added prior to infection. The macrophages were infected with Mtb at a multiplicity of infection (MOI) of 1 and incubated at 37°C with 5% CO_2_ for 3 hours, followed by treatment with 200 µg/ml amikacin for 1 hour. Amikacin was removed by washing infected monolayers with PBS twice and fresh medium containing relevant reagents was added before incubation for a further 24 hours. For CFU and MPN counts, the infected macrophages were washed with PBS and lysed with 0.2% (v/v) Triton X-100.

### Assessment of Viable Counts

MPN counts were quantified as described before.^3,7^ Briefly, sealed 48-well plates with serially diluted mycobacteria in 7H9 medium (MPN_7H9) or 7H9 medium supplemented with 50% (v/v) culture supernatant (MPN_CSN) were incubated at 37°C for up to 12 weeks. The MPN calculator program was used for determination of MPN counts.^58^ For CFU counts, 10 µl drops from each well corresponding to 10^-1^-10^-5^ dilutions from the MPN plates (7H9 and CSN) were spotted onto Middlebrook 7H10 agar plates supplemented with 10% (v/v) ADC and 0.5% (v/v) glycerol and incubated at 37°C for up to 12 weeks. In some experiments 7H10 agar was supplemented with 0.4% (w/v) charcoal; however, it did not increase CFU counts. Resuscitation Index (RI) was defined as RI=Log_10_ MPN/ml - Log_10_ CFU/ml. Limits of detection for CFU and MPN counts were 24 CFU/ml and 4.6 cells/ml, respectively. For assessment of viable counts LIVE/DEAD^TM^ *Bac*Light^TM^ Bacterial Viability kit (Thermo Fisher Scientific) and flow cytometry were used.

### Treatment of Mtb and BCG with NOD and CC

The NO donor 3-cyano-5-nitropyridin-2-yl diethyldithiocarbamate (NOD) and the control compound 3-cyano-4,6-dimethyl-5-nitropyridin-2-yl piperidine-1-carbodithioate (CC) were synthesized and charterised as described in the *Supplementary Material*. Purity of all compounds was >98%. Crystal structures of compounds were deposited in the Cambridge Crystallographic Data Centre (CCDC no. 2301117 and 2301118).

For treatment experiments Mtb or BCG was grown in 7H9 medium to OD_580nm_ of 0.8-1.0 at 37°C with shaking at 100 rpm. Cells were diluted with fresh 7H9 medium to an OD_580nm_ of 0.1 in a total volume of 5 ml. NOD or CC were dissolved in DMSO and added to cultures to a final concentration of 100 µM which corresponds to 30x the minimum inhibitory concentration (MIC) of NOD. Cultures were incubated at 37°C without shaking for up to 48 hours. Washing of NOD treated mycobacteria did not improve growth on agar or liquid media and was not used routinely.

### Detection of NO release in media and cells

NO release was estimated by the accumulation of its stable oxidation product, nitrite, using Griess reagent method.^26^ NOD or CC was incubated in 7H9 medium (pH 6.8) at a final concentration of 100 or 200 µM at 37°C without shaking for 24 hours. Absorbance at 540 nm was measured at different time points. Known concentrations of sodium nitrite were used to construct a calibration curve. For determination of NO in mycobacterial lysates and spent media, BCG cultures were grown to OD_580_∼0.8-1.0, centrifuged and resuspended in fresh medium (OD_580_ of 0.5). Independent cultures (10 ml) were set up in triplicate for each condition and treated with 200 µM NOD or CC or DMSO-only control (untreated control). At 1, 4 and 24 hours, 2 ml aliquots were taken and centrifuged at 13,000 x *g* for 5 minutes. The medium was removed and kept for nitrite determination; the pellets were washed with PBS and resuspended in 400 µl PBS containing Roche cOmplete™ Protease Inhibitor Cocktail and lysed by bead beating. Samples were centrifuged; supernatants and spent media were used for Griess Reagent System (Promega) by employing the internal standard provided by the kit. Intracellular concentrations of nitrite were normalised to protein content determined by Pierce™ BCA Protein Assay Kits.

For detection of NO in treated mycobacteria DAF-FM diacetate (4-Amino-5-Methylamino-2’,7’-Difluorofluorescein),^27^ purchased from Thermo Fisher Scientific, was used. BCG was pretreated with 10 µM of DAF-FM diacetate for 2 hours, followed by washing with 0.85% (w/v) NaCl or 7H9 medium three times before treatment with NOD and CC (100 µM). For flow cytometry BCG was incubated with either NOD or CC for 24 hours, washed with 0.85% (w/v) NaCl, stained with 10 µM of DAF-FM for 1 hour, followed by centrifugation and incubation in fresh 0.85% (w/v) NaCl for 15 minutes. DAF-FM diacetate-stained bacteria were analysed in CytoFLEX flow cytometer and CytExpert software (Beckman Coulter) or fluorescence was measured in Varioskan Flash using filters for excitation 495 nm and emission 515 nm.

### Live/dead staining and flow cytometry analysis

For assessment of viable counts LIVE/DEAD BacLight staining kit (Thermo Fisher Scientific) was used according to the manufacturer’s instructions. Stained BCG samples were analyzed using FACSAria flow cytometer and BD FACSAria^TM^ software (BD Bioscience). The following parameters were applied: SYTO 9 excitation/emission at 480/500 nm; Propidium Iodide excitation/emission at 490/635 nm. Ten thousand events were recorded from each sample; single parameter fluorescence overlay plots and dual parameter dot plots were generated. The side scatter-versus the forward scatter plot was generated from unstained samples and was used to set a gate for detection of bacteria. The side scatter-versus FITC-A dot plots (DAF-FM diacetate staining) or the FITC-A versus PE-Texas Red-A dot plots (LIVE/DEAD staining) were used to gate bacteria based on their fluorescence profile. The percentages of bacteria with distinct staining profiles were calculated. For each sample 10,000 events were recorded. Heat-killed BCG were included as a control.

### Transcriptomic analyses

RNA was extracted from three biological replicates of Mtb cultures after 4 hours of treatment with either NOD or CC using the GTC/Trizol method.^59^ After 4 hours of NOD treatment the bacteria retained culturability, thus enabling the study of adaptation during transition to the DC state. RNA (2 µg) was labeled with Cy3 and Cy5 fluorophores and hybridized to an Mtb microarray as previously described.^47^ Differentially expressed genes in NOD-compared to CC-treated cultures, 4 hours after exposure, were identified using a modified t-test (GeneSpring 14.5; Agilent Technologies) with Benjamini and Hochberg multiple testing correction and defined as those with p<5×10^-5^ and minimum fold change of 2.0. Microarray data have been deposited to the EBI BioStudies and are available under accession number is E-MTAB-10776.

For quantitative RT-PCR, RNA was reverse transcribed to cDNA using SuperScript II reverse transcriptase kit (Thermo Fisher Scientific) with mycobacterial genome-directed primers.^60^ The qPCRs were run in the *Corbett Rotor*-*Gene 6000* (Qiagen) using the 2×SYBR green master mix (Thermo Fisher Scientific) and primers (Table S1). Levels of expression were normalized to 16S rRNA. Relative gene expression in NOD-treated samples was calculated as the ratio of normalized gene copy number in NOD-treated samples to normalized gene copy number in CC-treated samples and expressed as relative Log_2_ fold change.

### Detection of Rpf peptides using mass-spectrometry

Mtb was grown in Sauton medium to OD_580_ 0.8. Filtered CSN was enriched on DEAE-Sepharose and digested with trypsin. LC-MS/MS was carried out using an RSLCnano HPLC system (Dionex, UK) and a Thermo Scientific LTQ Orbitrap Velos Mass Spectrometer. The raw data were processed using Proteome Discoverer (version 1.4.0.288, Thermo Scientific), Mascot (version 2.2.04, Matrix Science Ltd).

### Statistical analyses

Unpaired t-test or one-way ANOVA (Prism 10) were used to evaluate the statistic differences in the growth and flow cytometry datasets. Differentially expressed genes in ND-compared to CC-treated cultures, 4 hours after exposure, were identified using a modified t-test (GeneSpring 14.5; Agilent Technologies) with Benjamini and Hochberg multiple testing correction and defined as those with p<1×10^-5^ and minimum fold change of 2.0.

## ACKNOWLEDGMENTS

We acknowledge the Centre for Core Biotechnology Services at the University of Leicester for support with Containment Level 3 experiments, flow cytometry and mass-spectrometry analyses. We would like to thank Dr Kate Gould in the Bacterial Microarray Group at St. George’s University of London. We are grateful to Prof W.R. Jacobs (Albert Einstein College of Medicine) for providing the *Mycobacterium tuberculosis* H37Rv strain.

