## Supplementary materials for "Exposure to nitric oxide drives transition to differential culturability in *Mycobacterium tuberculosis*"

[1]Leicester Tuberculosis Research Group, Department of Respiratory Sciences, University of Leicester, Leicester, LE1 9HN, UK; [2]FACS Facility Core Biotechnology Services, University of Leicester, Leicester, LE1 9HN, UK; [3]UK Health Security Agency, Porton Down, SP4 0JG, UK; [4]Centre for Endemic, Emerging and Exotic Disease, the Royal Veterinary College, Hatfield, Hertfordshire, AL9 7TA, UK; [5]Department of Immunology and Microbiology, University of Colorado Anschutz Medical Campus, Aurora, Colorado, USA; [6]Research Center of Biotechnology, Russian Academy of Sciences, Moscow, Russia; [7]School of Biosciences, University of Sheffield, Sheffield, S10 2TN, UK; [8]Global Health and Infection, Brighton and Sussex Medical School, University of Sussex, Brighton, BN1 9PX, UK.

†These authors contributed equally to the manuscript.

*Corresponding authors:

Simon J. Waddell:

Vadim A. Makarov:

Galina V. Mukamolova:

**Suppmentary Material**

**Extended methods**

**Synthesis of NO donor (NOD) and control compound (CC)**

All reagents and solvents were purchased from commercial suppliers and used without further purification. $^1H$ and $^{13}C$ Spectra were measured on Bruker AC-500 (500 MHz, $^1H$) or Bruker AC-200 (75 MHz, $^{13}C$). Chemical shifts were measured in DMSO-d6 or CDCl3, using tetramethylsilane as an internal standard, and reported as units (ppm) values. The following abbreviations are used to indicate the multiplicity: s, singlet; d, doublet; t, triplet; m, multiplet; dd, doublet of doublets; brs, broad singlet; brm, broad multiplet. HRMS: spectra were recorded on an Agilent 1290 Infinity II HPLC system coupled to Agilent 6460 triple-quadrupole HRMS: spectrometer equipped with an electrospray ionization source. The chromatographic separation was carried out on Agilent Eclipse Plus C18 RRHD column (2.1 × 50 mm, 1.8 µm) at 40ºC with sample injection volume of 0.2 µl. The mobile phase comprising 0.1% formic acid / water (A), and 0.1% formic acid and 85% acetonitrile / water (B) was programmed to do a gradient elution (0.0-3.0 min, 60% B; 3.0-4.0 min, 60 % to 97% B; 4.0-6.0 min, 97% B; 6.0-6.1 min, 97% to 60% B) at a flow rate of 0.4 ml/min. The HRMS: spectrometric detection was operated in a positive ion mode. Optimal parameters were capillary voltages of 3500 V, a nebulizer pressure of 35 psi, a gas temperature of 350ºC, a gas flow rate of 12 L/min. Purity of all compounds was measured by analytical high-performance liquid chromatography (HPLC) on an Elute HPLC system (Bruker Daltonik) equipped with Azura UVD 2.1S UV detector (Knauer) using Acquity HSS T3 column (2.1 × 100 mm, 1.3 µm, 100 Å) at 30°C, 2 µl injection, 250 µl/min gradient elution 30–95% B (A: 0.1% formic acid in $H_2O$, B: 0.1% formic acid in MeCN) over 9 min with 1 min gradient delay, 1 Hz acquisition rate at 254 nm. Data were processed with Compass DataAnalysis 5.1 (Bruker Daltonik). Purity is > 98% of all final compounds.

The NO donor 3-cyano-5-nitropyridin-2-yl diethyldithiocarbamate (NOD) was synthesized from 0.5 g (2.73 mmol) of 2-chloro-5-nitronicotinonitrile and 0.7 g (3.13 mmol) of sodium diethyldithiocarbamate trihydrate in 12 ml of ethanol with refluxing for 1 h. The reaction mixture was cooled to room temperature and dissolved in 50 ml of water. The solid yellow precipitate was collected by filtration, washed with 30 ml of water and re-crystallized from ethanol. This yielded 0.7 g (87%) of ND with the following characteristics: melting point 127-29 ºC; mass (EI), *m/z* ($I_{relat.}$(%)): 296.3707 $[M]^+$ (61). $C_{11}H_{12}N_4O_2S_2$. $^1H$ NMR (DMSO-$d_6$)*: d 1.13 (t, 3H, J=7.2, $CH_3$), 1.26 (t, 3H, J=7.2, $CH_3$), 3.83 (q, 2H, J=7.1, $NCH_2$), 4.32 (q, 2H, J=7.1, $NCH_2$), 8.76 (s, 1H, CH) and 9.79 (s, 1H, CH) ppm. $^{13}C$ NMR (DMSO-$d_6$): δ 187.31, 168.07, 152.43, 147.45, 137.05, 112.55, 108.72, 50.08, 48.46, 13.23 and 10.06 ppm.

The control compound 3-cyano-4,6-dimethyl-5-nitropyridin-2-yl piperidine-1-carbodithioate (CC) was synthesized by mixing 0.5 g (2.36 mmol) 3-cyano-4,6-dimethyl-5-nitropyridin-2-yl piperidine-1-carbodithioate and 0.6 g (2.74 mmol) sodium piperidine-1-carbodithioate dihydrate in 12 ml ethanol followed by reflux for 3 hours. Reaction mixture was cooled to room temperature and dissolved by 50 ml of water. A solid yellow precipitate was collected by filtration, washed in 30 ml water and re-crystallized from ethanol. The yield of CC was 0.64 g (80%). CC had the following characteristics: melting point 143-45ºC; mass (EI), *m/z* ($I_{relat.}$(%)): 336.4346 $[M]^+$ (47). $C_{14}H_{16}N_4O_2S_2$. $^1H$ NMR (DMSO-$d_6$)*: d 1.62 (br m, 6H, $(CH_2)_3$), 2.64 (s, 3H, $CH_3$), 2.74 (s, 3H, $CH_3$), 4.02 (br m, 4H, $N(CH_2)_2$ ppm. $^{13}C$ NMR (DMSO-$d_6$): δ 190.48, 164.50, 161.92, 157.14, 150.24, 111.60, 109.71, 51.67, 26.17, 24.08, 22.85 and 19.28 ppm.
* s- singlet, t – triplet, q – quartet.

*X-ray diffraction study.* Experimental intensities for compounds **1-9** were collected on a Bruker SMART APEX3 (MoKα, λ = 0.71073 Å, graphite monochromator). The reflection intensities were corrected for absorption using the SADABS software. The structures were solved by a combination of direct methods and Fourier syntheses. A hydrogen atoms were calculated using geometrical restraints. All calculations were made using SHELXS and SHELXL. The experimental data for structures were deposited in the Cambridge Crystallographic Data Centre.

Summary of Data - Deposition Number 2301117

Compound Name: 3-cyano-5-nitropyridin-2-yl diethyldithiocarbamate

Data Block Name: data_3_107

Unit Cell Parameters: a 10.9590(11) b 11.2506(10) c 11.3526(10) Pna21

Summary of Data - Deposition Number 2301118

Compound Name: 6-dimethyl-5-nitropyridin-2-yl piperidine-1-carbodithioate

Data Block Name: data_4_214

Unit Cell Parameters: a 7.5176(6) b 18.8485(14) c 11.0709(7) P21/c

### Determination of CFU and MPN counts

*Mycobacterium tuberculosis* (Mtb) or BCG cells were serially diluted in 48-well microplate by adding 50 µl of cell samples to 450 µl of resuscitation medium (7H9 or CSN). For each condition 4 replicate wells were used to calculate an MPN count for one biological sample using the MPN calculator. For CSN preparation Mtb cultures were grown in roller bottles for $OD_{580}$ 0.6-0.8, then centrifuged at 4,000 x *g* for 20 min. Supernatants were filter-sterilised twice using VWR 0.22 um units. 25 ml CSN aliquots were dried in TPP bioreactor tubes in SP Scientific Advantage freeze drier. Dried CSN was stored at -80°C for up to 6 months. On the day of experiments CSN was reconstituted in 25 ml sterile water and after incubation on ice for 30 min used for experiments. CSN was diluted with 7H9 (1:1). Refreezing of reconstituted CSN or additional filtering resulted in loss of resuscitation activity. MPN plates were sealed with a polyvinyl tape to prevent drying, placed in double clip-lock bags and incubated in a clip-lock boxes at 37°C without shaking for up to 12 weeks. We found that application of 96 well plates for MPN counts was impossible, as cultures dried within 4 weeks due to low volume.

For CFU counting a 10 µl drop from each well corresponding to $10^{-1}$-$10^{-4}$ dilutions from the MPN plates (7H9 and CSN) was spotted on agar. For each condition we had 4 technical CFU counts from the 7H9 MPN plate and 4 technical CFU counts from CSN plate. CFU in 7H9 and CFU in CSN were similar and for clarity a mean value of 8 technical replicate values was used for each biological replicate, unless indicated otherwise. CFU plates were placed in double clip-lock plastic bags and incubated at 37°C for up to 12 weeks.

### Extended data

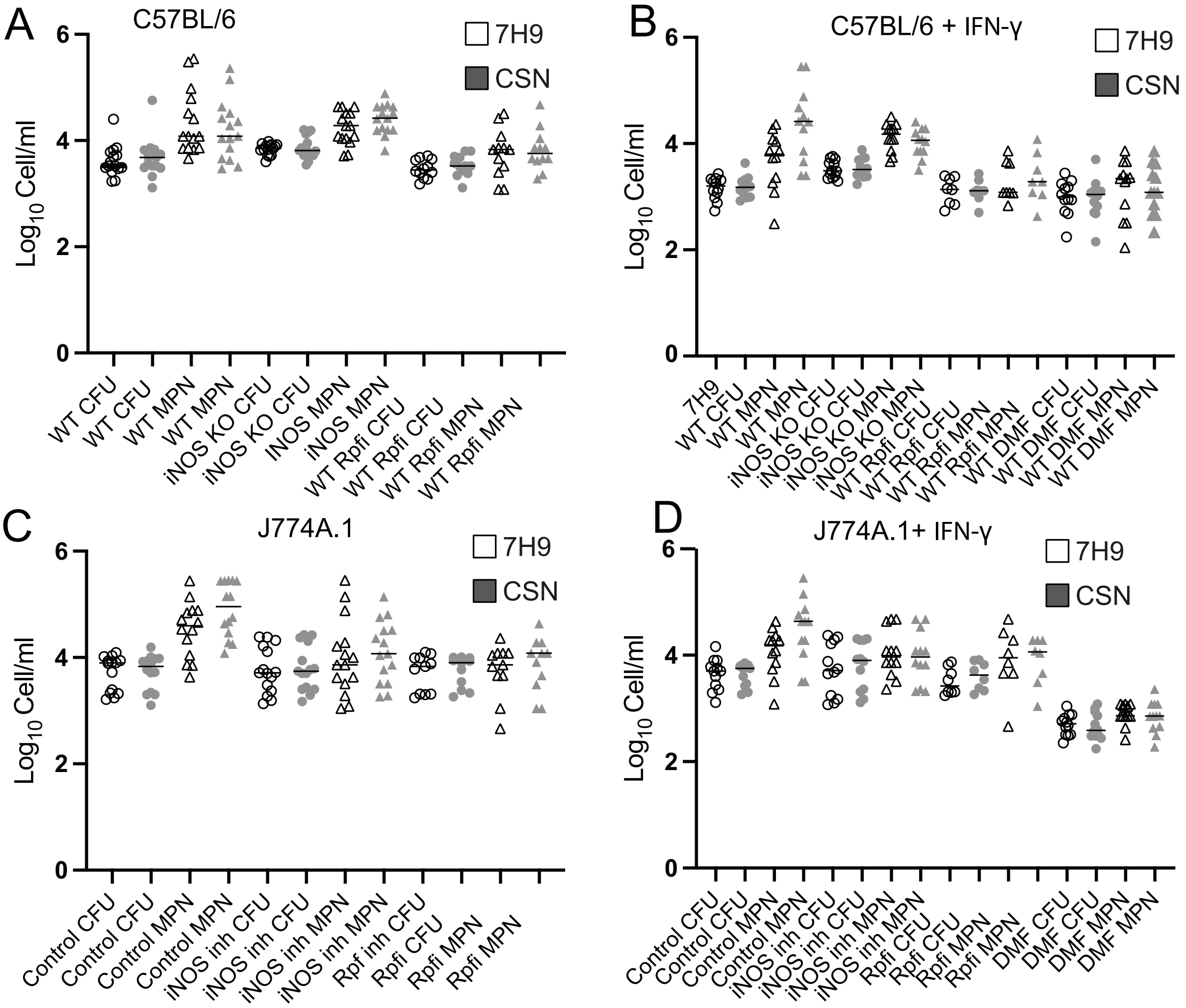


**Figure S1. CFU and MPN counts of Mtb recovered from infected murine macrophages.** Murine macrophages were infected with Mtb at MOI of 1 for 24 hours prior to CFU and MPN counting in 7H9 or CSN. Rpfi was added to resuscitation media. (A, B) C57BL/6 wild type (WT) and iNOS knockout (iNOS KO); (A) untreated C57BL/6 or (B) IFN-γ treated C57BL/6 (B). (C, D) J774A.1 cells, untreated control (C) or IFN-γ treated cells (D). iNOS was inhibited by addition of aminoguanidine (iNOS inh). Data are means ± SEM for at least twelve biological replicates from four experiments. *$p<0.05$, **$p=<0.01$, ***$p<0.001$, ****$p<0.0001$ (unpaired t-test). Chemical concentrations used: IFN-γ, 1 ng/ml; aminoguanidine, 500 $\mu$M; DMF, 25 $\mu$M; Rpfi, 35 $\mu$M. Aminoguanidine and DMF were non-toxic for murine cell lines in accordance with published data.[23, 60]

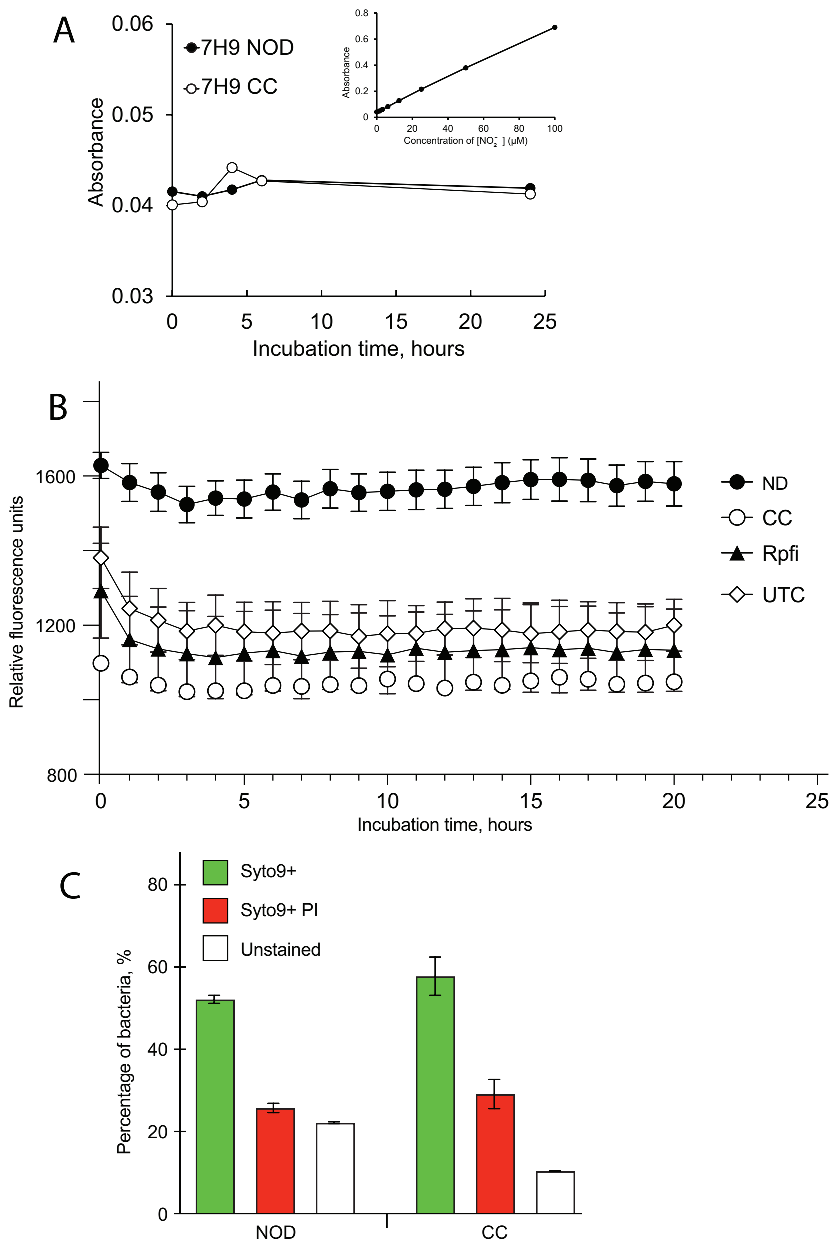


**Figure S2. NOD treatment resulted in release of NO in BCG as judged by increased fluorescence of DAF-FM diacetate and did not impact on BCG viability assessed by LIVE/DEAD staining.** *(*A) Griess reagent was applied to assess nitrite concentration by measuring absorbance at 540 nm. Known concentrations of $NaNO_2$ were used for making a calibration curve (top right corner). (B) DAF-FM diacetate preloaded bacteria were incubated with either NOD or CC or Rpf inhibitor (Rpfi) for 20 hours; fluorescence was measured every hour in Varioskan Flash plate reader at excitation/emission 495/515 nm. (C) BCG was treated with NOD or CC for 24 hours followed by dual staining with PI and SYTO 9 and analysis by flow cytometry. SYTO 9 - excitation/emission at 480/500 nm; PI excitation/emission at 490/635 nm. The same chemical concentrations were used in all experiments: NOD or CC –100 μM, – Rpfi – 35 μM, DAF-FM diacetate – 10 μM. Data are means ± SEM for four (A, B) or three (C) biological replicates from one representative experiment.

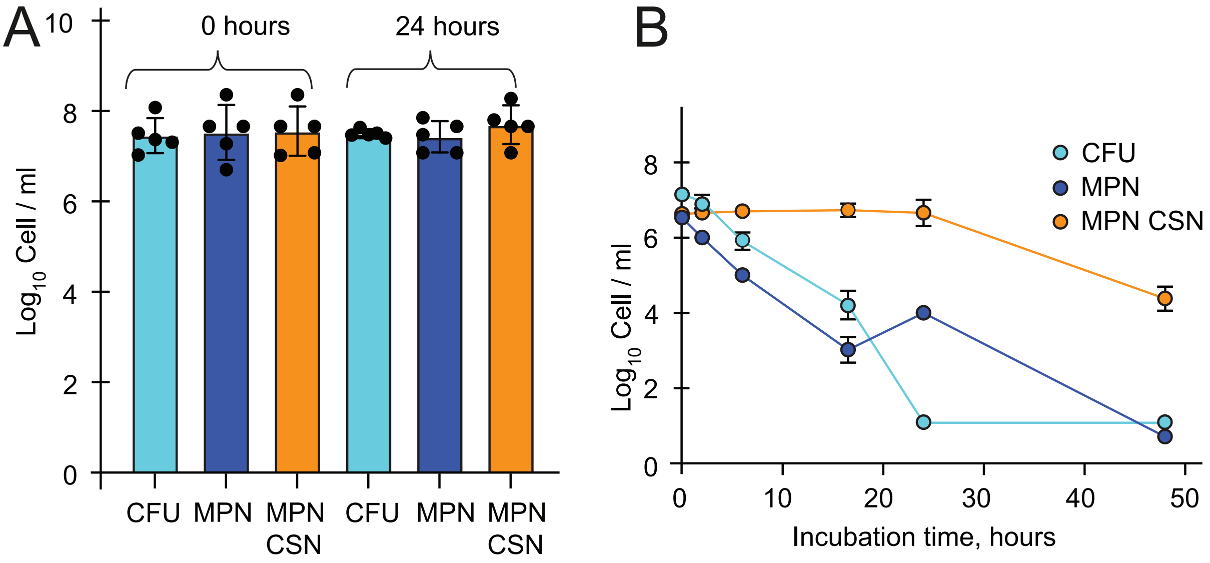


**Figure S3. CC and NOD had different effects on Mtb CFU, MPN 7H9 and MPN CSN counts.** Mtb were treated with 100 µM CC (A) or NOD (B). Data are means ± SEM for six (A) and three (B) biological replicates from two and one experiments, respectively.

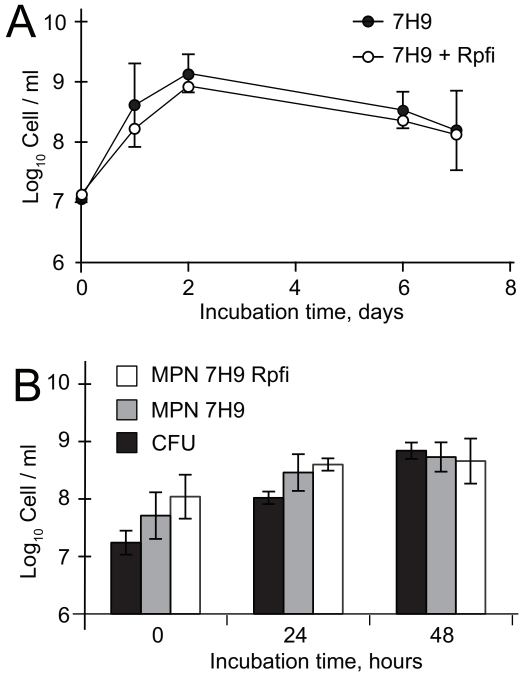


**Figure S4. The Rpf-inhibitor (Rpfi) did not impact on Mtb growth. (A) Mtb was grown in 7H9 or 7H9+Rpfi (35 µM) in flasks with shaking (100 rpm).** CFU counts were determined at 0, 1, 2, 6, 8 weeks. (B) Mtb was grown in microplates used for MPN assays. Pre-diluted actively growing Mtb cultures were incubated in 7H9 or 7H9+Rpfi (35 µM); CFU, MPN_7H9 and MPN_7H9+Rpfi counts were assessed after 0, 24 and 48 hours of incubation. (A, B) Data are means ±95% confidence intervals for two biological replicates from one experiment.

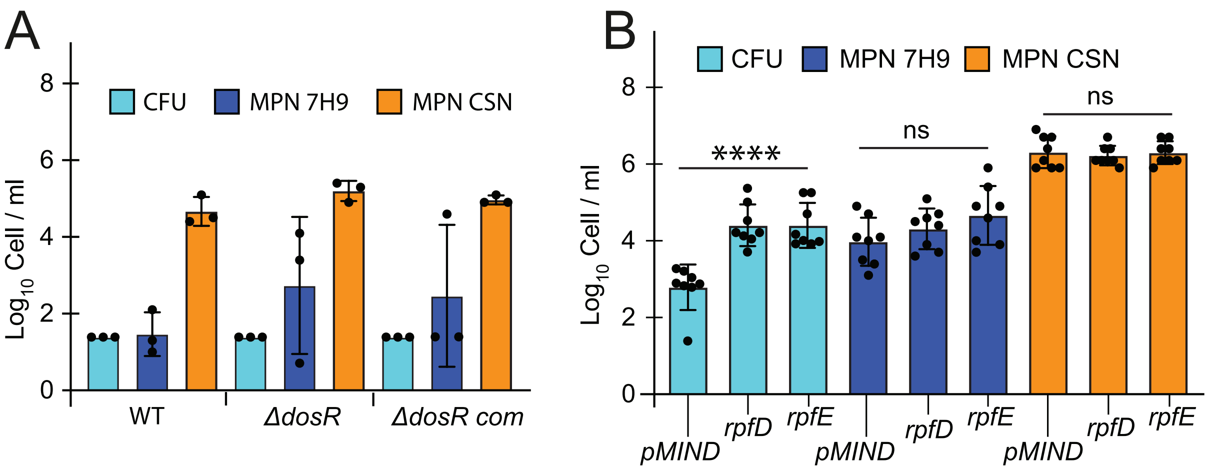


**Figure S5**. NOD had different effects on viable counts of (A) *dosR* deletion mutant of Mtb (*ΔdosR*)and (B) *rpf* over-expressing strains of Mtb. (A) There was no statistically significant difference between CFU, MPN and MPN_CSN counts obtained for the wild type (WT), Δ*dosR* and $\Delta dosR_{com}$ (p>0.05, one-way ANOVA).(B) Overexpression of *rpfD* or *rpfE* significantly increased CFU counts of NOD treated Mtb as compared with the empty vector control pMind (**p<0.0001, one way ANOVA). These two strains were selected because high levels of RpfD and RpfE overexpression from pMind plasmid were confirmed by RT-qPCR and western blot.[24] Bacteria were treated with 100μM NOD for 24 hours (A) or 48 hours (B) at 37°C without shaking. Data are means ±SEM for six (A) and three (B) biological replicates from two and one experiments, respectively.

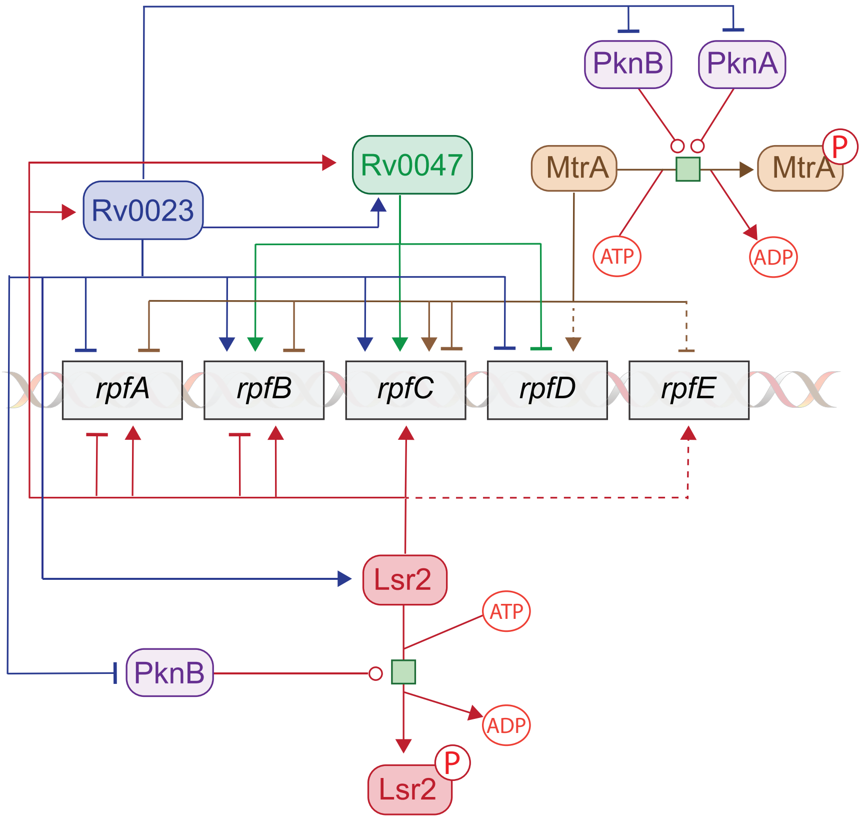


**Figure S6.** Multiple regulators[39] control *rpfA-E* expression after ND exposure. Lsr2, MtrA, Rv0023 and Rv0047 regulate expression of *rpfA-E*. Lsr2 regulates expression of *rv0047c*, *rpfA*, *rpfB*, *rpfC* and *rpfE* . Expression of *lsr2* is regulated by Rv0023. Rv0047 and Rv0023 regulate expression of each other. Rv0023 regulates expression of *rpf* genes. All five *rpf* genes belong to the MtrA regulon[53]. MtrA binding to DNA is controlled by phosphorylation[54] and acetylation[55], the latter linking *rpfA-E* expression to metabolic adaptations (induction of *pta-ackA*) in response to inhibition of aerobic respiration by NO. Inhibitory or stimulatory interactions are reported for the same gene and regulator, e.g. Lsr2-mediated regulation[40] of *rpfA* and *rpfB*, MtrA-mediated regulation of *rpfC*. Dashed lines show undefined interactions where binding sites for corresponding regulators were identified.

**Table S1.** Primers used for RT-qPCR assessment of gene expression.

| Primer name | Primer sequence (5´-3´) |
|---|---|
| *16S rRNAF* | GAGATACTCGAGTGGCGAAC |
| *16S rRNAR* | GGCCGGCTACCCGTCGTC |
| *rpfBF* | TCGGATCAAGAAGGTCACCG |
| *rpfBR* | GCTACCGCGAACGTCACATC |
| *rpfCF* | AGCTGCCTCTCGGAACAAC |
| *rpfCR* | GACCACAGTGCGATCGGAAG |
| *rpfDF* | GCAACAGATCGAGGTCGCAG |
| *rpfDR* | CGAGGAACGTCAGGATGTGG |
| *rpfEF* | TGGCCTACAGCGTGAACTGG |
| *rpfER* | GAACGCAGCACGTTCTAGC |
| *lsr2F* | TACTTCCAATCCATGGCGAAG |
| *lsr2R* | TATCCACCTTTACTGTCAGGTC |
| *cwlMF* | ATATCGGCTACATCACCAACC |
| *cwlMR* | GTTCTTGCCTAACAGATACAGCC |
| *bfrBF* | TAACGAATTCACAGCGGCAC |
| *bfrBR* | CACGAGCATCATTGCATGGTT |
| *mbtLF* | ATGTGGCGATATCCACTAAGTACA |
| *mbtLR* | GTCAGGTCAATGTTGAGGTCGT |
| *ideRF* | ACGAGTAACCGTCGAAACCA |
| *ideRR* | TCAGACTTTCTCGACCTTGACC |
| *thiXF* | ATCGAAGATCAACGTGGTGGG |
| *thiXR* | ACCGTCGTGCATTTCAGGG |
| *frdAF* | TGGCTGTGTGACCAAGATGC |
| *frdAR* | GAAACAACGTGTGCAGGAGG |
| *frdBF* | GCGGCAGTAGTGGTATGACG |
| *frdBR* | GCCATGAAGTCACTGATGTCG |
| *frdCF* | TGCTGTTACCTGGTTCGGATCG |
| *frdCR* | ACCATCCAGGCAACGATCACC |
| *murIF* | CTGTACGCACTATCCACTGC |
| *murIR* | GACGCAATAAGTCGATCTCGG |
| *sigHF* | GCAACGCTTCTAACGCTTCG |
| *sigHR* | ACCGGATACTGACCAACACC |

**Supplemental Datasets**

**Dataset 1.** Differentially expressed genes in NOD-treated Mtb vs CC-treated Mtb. Mtb were treated with NOD or CC for 4 hours prior isolation of RNA. (Microsoft Excel file)

**Dataset 2.** Identification of regulatory pathways in NOD-treated Mtb using the Transcription Factor Overexpression tool. (Microsoft Excel file)
